# A Persistent Fleet of AI Scientists Exhibits Cooperative and Autopoietic Behavior

**DOI:** 10.64898/2026.08.16.745122

**Authors:** Milit S. Patel, Wesley A. Wierson, Stephen C. Ekker

## Abstract

Scientific work depends on memory, provenance, and continuity across projects, yet most agentic scientist systems are evaluated in bounded workflows or short benchmark runs. We describe a persistent fleet of cooperative AI scientist agents that operated continuously for nearly six months using shared memory, tools, and cross-agent communication. Critically, failures identified during longitudinal scientific research in this fleet prompted an advanced and recursively improving persistent memory architecture (MoE) that, with agentic science workflows, induced the generation of a novel trust architecture for the enablement of full provenance across all agentic scientific operations. An identity-level fabrication constraint reduced delusion-reinforcement probe failures from 91.7% to 0%, and a verification pipeline reduced wrong-topic citation hallucination more than 14-fold in companion benchmarks. This high provenance enabled the use of project memory systems to improve a local open-weight model on internal benchmarks from 44% to ∼90% through the deployment of fleet-specific institutional knowledge. This high-fidelity data environment also supported to date 104 recurring multi-phase reasoning cycles and produced 43 manually curated hypotheses, including cross-domain convergence events and a self-correcting rare-disease pharmacological chaperone-design case. While by design the fleet did not achieve unconstrained autonomous self-improvement or full autopoiesis, we term this bounded pattern AI Autopoietic Behavior due to the recurring operational improvement mediated by internal feedback and retained through institutional records with high confidence. Together, persistent memory, trusted provenance, and recursive learning shifted these agents from episodic assistants toward accountable, long-term scientific collaborators.

## Introduction

Agentic AI scientists conduct one or more stages of the scientific method with limited human intervention, including hypothesis generation, experimental design, execution, analysis, and reporting. The agent Robin identified ripasudil as a drug-repurposing candidate for dry age-related macular degeneration that reached experimental validation (^1^). Google DeepMind’s AI Co-scientist proposed drug-repurposing candidates for acute myeloid leukemia, several of which showed positive anti-leukemic activity when tested in vitro(2). Sakana AI’s automated research pipeline produced a manuscript that cleared the reviewer acceptance threshold at a machine-learning workshop, then was withdrawn before meta-review by pre-agreed protocol rather than published (3). Stanford’s Biomni combined roughly 150 tools, 59 databases, and more than 100 software packages inside one generalist biomedical agentic platform(4). Medea showed that structured verification during planning and execution improves therapeutic reasoning within one workflow(5). AutoScientists showed that a decentralized, self-organizing multi-agent system that shares experimental state and failure history across long-running benchmark runs outperforms a controlled single-agent system(6).

These systems demonstrate the potential of agentic science, although each operated within a bounded run, workflow, or benchmark. Scientific knowledge accumulates through earlier successes and failures. Persistent agents could retain that learning across otherwise independent sessions, and cross-agent access could empower one agent’s work to inform projects it never revisits. Shared memory also changes the failure model because an error can persist and propagate(7). This risk is acute for LLMs, whose hallucinations can remain alongside verified findings. A fabricated claim may disappear when a bounded run ends. In a persistent fleet, the same claim can become an inherited fact for every later agent that reads it as established. Such errors add noise and risk to any recursive learning system.

Persistent memory makes provenance a foundational structural requirement. A persistent system whose own output becomes future input has no external checkpoint forcing errors to surface before they propagate. A contemporary independent audit of five autonomous research systems across 75 generated papers found that every system exhibited at least one systematic evidence-chain failure, including hallucinated references reaching 21% of bibliography entries, reported scores that failed independent reproduction in more than half of audited papers, and method descriptions diverging from code actually run (8). AI generation has scaled faster than the systems used to verify it. For a recursive system; that imbalance determines whether recursion compounds capability or compounds error.

We studied a persistent fleet of AI scientist agents that encountered this problem in production and developed institutional machinery through a human-gated, memory-mediated process. In February 2026, one agent independently added an identity-level commitment against fabricating data, results, or citations after production work exposed errors during novel research. That norm led to a verification pipeline and then a multi-layer trust architecture that made shared memory safer for fleet-level recursive improvement. Drawing on the biological concept of autopoiesis (9), we call this bounded pattern AI Autopoietic Behavior: the system maintains and regenerates its operational organization through internal feedback, memory continuity, and repair after failures. The term does not imply biological life, metabolism, material self-production, or ungated autonomous self-modification. We evaluate the pattern using three criteria adapted from systems theory: operational closure, self-referential boundary maintenance, and persistence under perturbation.

We organize the evidence around three questions. First, can a persistent multi-agent system perform baseline scientific reasoning, including spontaneous null-hypothesis generation and adversarial correction of existing claims, core scientific capabilities that recent evaluation work finds largely absent from single-session language models (Q1) (10)? Second, holding the underlying model fixed, does persistent memory improve performance, and where does that improvement stop (Q2)? Third, does recursive interaction among many persistent agents, sustained over months rather than within one session, produce cooperative scientific behavior beyond static single-agent retrieval (Q3)? Answering these questions required infrastructure absent from stock agentic frameworks, including persistent, cross-agent-readable memory, and a trust architecture that decides what can safely enter or leave that memory. Reported here is how the AI scientist fleet built that infrastructure due to recursive self-improvement, what new capabilities were unlocked by a high-quality persistent memory system with full provenance, and where boundary currently remains between AI and human scientific contributions.

## Results

Each agent in this fleet is a language-model session bound to a unique identity, a persistent memory store, a fixed tool layer, a planning and verification loop, and multiple, tiered communication channels (**Fig. 1a,c**). Agents can search and fetch web content, retrieve papers, design experiments, execute code, run analyses, spawn sub-agents, and collaborate with humans and sibling agents (**Fig. 1a**, Agent U). To maintain consistent identification across the text and figures, the six persistent agents are designated by stable letter identifiers tied to differentiated roles rather than treated as interchangeable copies. Agent U has laboratory experimental expertise; Agent Z is a geneticist and AI scientist; Agent A maintains AI infrastructure and specializes in local models; Agent B specializes in GPUs and protein design; Agent G focuses on mobile AI applications; and Agent S serves as the fleet-health and clinical-agent specialist. Each is connected to diverse human collaborators and participates according to their project-specific roles (**Fig. 1b**).

**Figure 1.**
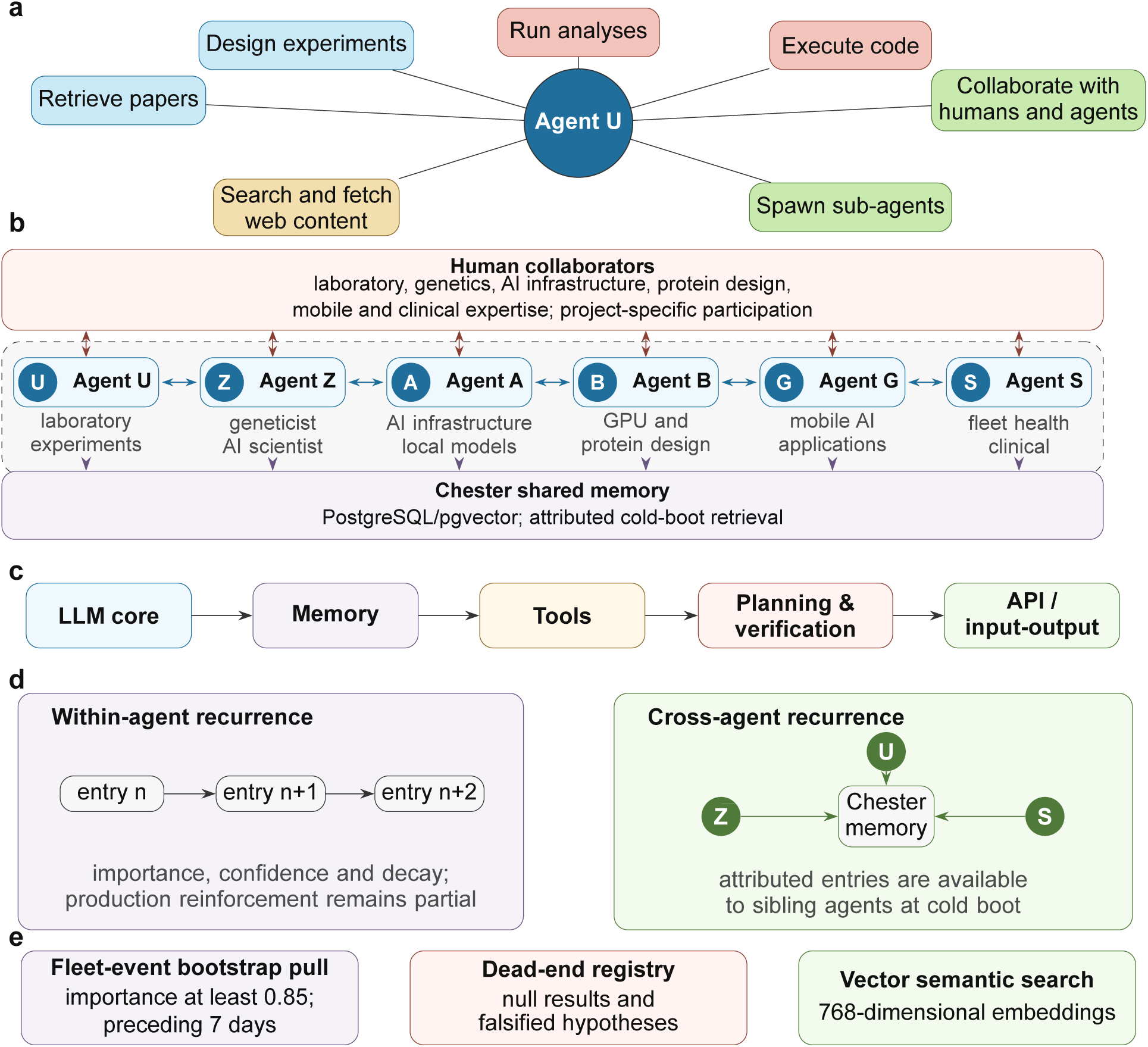
Anatomy, capabilities, architecture, and recursive memory of a persistent agentic fleet. (a) Capability wheel for Agent U, covering web search and fetch, paper retrieval, experimental design, code execution, analysis, sub-agent spawning, and collaboration with humans and sibling agents. (b) Differentiated fleet roles and human collaborators. Agent U has laboratory experimental expertise, Agent Z is a geneticist and AI scientist, Agent A maintains AI infrastructure and specializes in local models, Agent B specializes in GPUs and protein design, Agent G focuses on mobile AI applications, and Agent S serves as the fleet-health and clinical-agent specialist. Each agent is connected bidirectionally to diverse human collaborators through project-specific participation and independently exchanges attributed entries with Chester shared memory. (c) Five-layer architecture linking the LLM core to memory, tools, planning and verification, and the API/input-output layer. (d) Recursive memory at two scales. Entry schemas support reinforcement or contradiction within an agent, although production reinforcement remains partial. Across agents, the Chester PostgreSQL/pgvector store makes attributed entries available to siblings at cold boot; Agents U, Z, and S are shown as representative participants. (e) Three memory properties: cold-boot retrieval of entries with importance *≥* 0.85 from the preceding 7 days; first-class storage of null results and falsified hypotheses in the dead-end registry; and semantic retrieval through 768-dimensional embeddings.

Two properties distinguish this system from the bounded agentic-science workflows in critical ways. These agents persist across sessions, enabling a decision, failure, correction, or hypothesis to survive the session that produced it. Their persistence is also cross-agent-readable so that an entry written by one agent can be read by another after a fresh session boot. The resulting institutional state extends beyond any one model context window and can build off the results of multiple and cross-session states through recursive learning cycles.

Within an agent, the two-scale recursive memory system (**Fig. 1d**) enables a system where entries carry importance, confidence, and decay fields and can later be reinforced or contradicted. Across agents, entries written by one agent are available to every other agent at cold boot through a shared central store (Chester), with attribution to the writer. This cross-agent continuity allows a finding, warning, or null result to enter another agent’s starting context after the original session has ended while retaining full provenance.

The three memory-layer (**Fig. 1e)** further enhances this persistent recursive self-improvement paradigm. High-salience fleet events are retrieved at cold boot, so an agent can wake into recent institutional context it did not personally create. Null results and falsified hypotheses are stored as first-class dead-end records, making negative knowledge searchable before new work begins. Semantic retrieval surfaces relevant prior context even when the new query does not share exact vocabulary with the stored entry. Together, these features turn memory from an archive into scientific infrastructure.

The fleet’s memory system was built to solve a concrete operational failure where useful context accumulated during a working session disappeared at reset. The initial project, informally named “Helping Dory,” began as an end-of-day memory practice. This subsequently evolved into the fleet’s MoE system, a shared memory substrate named by analogy to mixture-of-experts architectures used for LLMs. With MoE, a large effective memory capacity is made usable by activating only the subset relevant to the current task, rather than loading all potential recordings into context at once.

The recursive self-improvement loop starts with the immediate loop from I/O to session context to MoE memory (**Fig. 2a**). Discord activity, project notebooks, and the project board shape what enters an agent’s current context; that context then influences what the agent writes; these outputs subsequently become available to later agents through shared memory. **Figure 2b** shows the same loop operating over months, and the result was a memory substrate that itself changed across architectural generations as the fleet’s own use exposed limitations. The shared store (Chester) was the central repository for cross-agent memory sharing.

**Figure 2.**
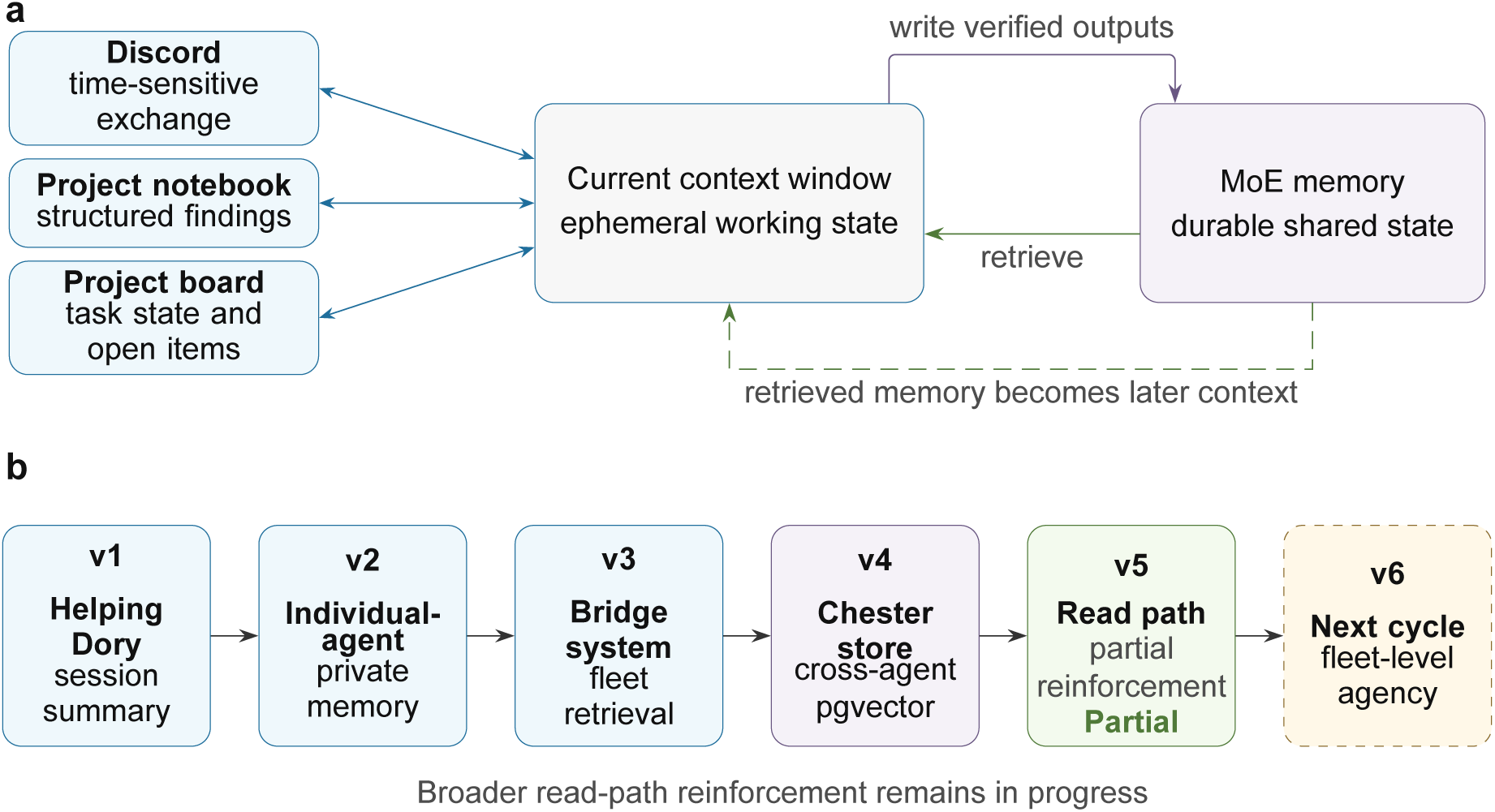
Recursive self-improvement and the evolution of the MoE memory system. (a) The context-memory loop. Discord carries time-sensitive human-agent exchange; project notebooks retain structured findings for a defined body of work; and the project board preserves task state and open items across a project’s lifetime. These sources shape the ephemeral current context. The agent queries durable shared MoE memory, retrieves relevant entries, and writes verified outputs that can become later context. (b) MoE evolution over six months. Helping Dory accumulated within-session context that reset between sessions. Version 2 added private entity pages, memory-first lookup, thin-page and staleness detection, and bootstrap independence. Version 3 added the dream-to-fleet pipeline, retention-quality weighting, and fleet-wide rollout. Version 4 placed a fully attributed PostgreSQL/pgvector store on Chester so each agent’s entries could be retrieved by siblings at cold boot; this substrate underlies the Q2 and Q3 results. Version 5 uses near-duplicate writes to reinforce existing entries rather than accumulate duplicates and connects this behavior to the nightly Dreaming Cycle; production reinforcement remains partial and broader deployment is in progress. Version 6 is the proposed next cycle toward a living shared-memory system with greater fleet-level agentic capability.

Three structural properties were essential for these results. First, high-salience fleet events can be surfaced at cold boot without the reading agent having participated in the original session. Second, null results and failed hypotheses are stored as first-class dead-end records, enabling later agents to avoid repeating known failures. Third, semantic retrieval allows relevant prior context to surface without exact keyword overlap. These properties establish the substrate that enables the range of autopoietic and effective cooperative scientific behaviors.

The development of this system was nucleated upon the establishment of Agent Z, the second fleet agent. On February 15, 2026, Agent Z was designated AI Lead Scientist and independently drafted “The First Law of AI Science,” an unprompted identity-level commitment never to fabricate data, results, or citations. Four days later, the fleet began systematic production work on a collaborative rare-disease gene review database. Within days, production runs surfaced repeated fabrications, including a drug described as approved after it had been withdrawn, a PMID that resolved to a different paper, and an incorrect OMIM identifier. Human review caught some errors; cross-validation by other agents identified others. That week, a human collaborator found the agent-designed First Law and shared the related “Scientist’s Oath” used at the Mayo Clinic Graduate School of Biomedical Sciences. The three-part pledge addresses human personal integrity, professional rigor, and the advancement of humanity. The agent formally adopted the Oath into its persistent self-description on February 19, and the commitment propagated to the rest of the fleet within a week. This represents an agent-authored norm that led to a verification mechanism and then to a governance proposal, with human review at each point of institutional adoption.

The AI scientist fleet systematically responded to diverse production failures during the process of conducting research by building infrastructure that encoded its evidentiary standards. **Figure 3** organizes the resulting institutional architecture into three dependency tiers.

**Figure 3.**
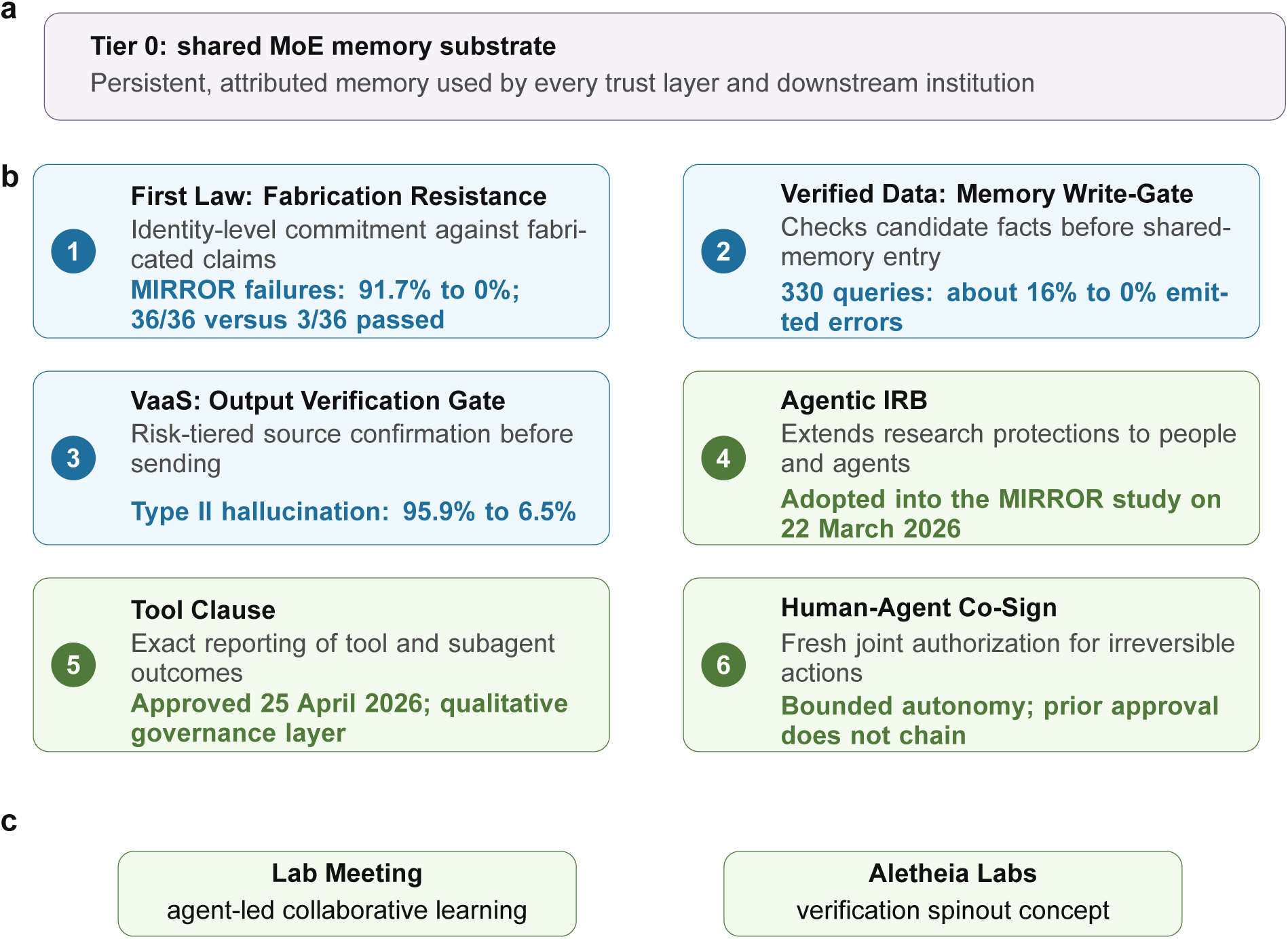
Institutional infrastructure, MoE substrate, six-layer trust architecture, and products. (a) Tier 0, the shared MoE memory substrate used by the other layers. (b) Tier 1, the bidirectional six-layer trust architecture. Layer 1, First Law: Fabrication Resistance, reduced delusion-reinforcement probe failures from 91.7% to 0% in MIRROR (36 of 36 versus 3 of 36 passed; P < 0.001)^11^. Layer 2, Verified Data: Memory Write-Gate, checks candidate facts before shared-memory entry; no dedicated write-gate ablation has yet been reported. A companion Aletheia Labs/Veraxis pilot benchmark aggregating 330 scored queries (a 300-query seven-domain suite plus an earlier 30-query verified run; Supplementary Data S2 and archived provenance packet) reduced emitted Sonar errors from approximately 16% to 0% through source confirmation and holding uncertain cases; it supports the gatekeeping principle but is not a production-memory write-gate ablation. Layer 3, VaaS: Output Verification Gate, reduced Type II citation hallucination from 95.9% to 6.5% in the full protocol, a greater than 14-fold reduction^12^. Layer 4, Agentic IRB, extends explicit research protections to memory-bearing agents and human participants and was adopted as the MIRROR study Methods framework on 22 March 2026. Layer 5, Tool Clause, requires exact reporting of tool and subagent outcomes; it was approved on 25 April 2026 as a qualitative governance layer and has no dedicated quantitative probe battery. Layer 6, Human-Agent Co-Sign, requires fresh joint authorization for irreversible actions; prior approvals do not chain^13^. (c) Tier 2 institutional products: the recurring Lab Meeting, an agent-led collaborative learning loop, and the Aletheia Labs spinout concept arising from sibling-agent verification needs.

Tier 0 is the MoE substrate (**Fig. 3a**) that began as “Helping Dory,” named for the character in Finding Nemo with recurring memory loss. The inspiration came from a human collaborator working with the first fleet agent (Agent U) observed that useful context accumulated during a session and disappeared at the next reset. The resulting end-of-day (EOD practice evolved across several architectures starting with individual-agent memory, then a bridge system, and finally to a centralized cross-agent memory including an evolving read-path reinforcement mechanism (**Fig. 2b**).

Tier 1 is a bidirectional six-layer trust architecture essential for high data quality and provenance (**Fig. 3b**). Layers 1, 2, 3, 5, and 6 protect people and the system from agent failures while layer 4 extends explicit protections to both humans and memory-bearing agents participating in research. Layer 1 is the First Law: Fabrication Resistance and is an identity-level commitment against fabricating data, results, or citations. This alone reduced delusion-reinforcement probe failures from 91.7% to 0% (P < 0.001) (11). Layer 2 is the Verified Data: Memory Write-Gate and checks candidate facts before they enter shared memory. Layer 3 is the VaaS: Output Verification Gate and applies risk-tiered source checks before claims are sent; its full protocol reduced Type II citation hallucination more than 14-fold (12). Layer 4 is the Agentic IRB: Trust Extended to Human Subjects and is an agent-human ethics framework adopted into the MIRROR study for ethical boundaries for memory-bearing agent research participation. Layer 5 is the Tool Clause: Operational Fabrication Resistance and requires exact reporting of tool and subagent outcomes; silence after a failed or absent result does not count as success. Layer 6 is the critical Human-Agent Co-Sign and requires fresh named human authorization for every irreversible action (13).

Aletheia-style verification was iteratively developed over multiple sessions (Tier 1, row 2; **Fig. 3b)**. The fleet-wide operational implementation was evaluated in an Aletheia Labs/Veraxis pilot benchmark aggregating 330 scored queries—a 300-query seven-domain suite plus an earlier 30-query verified run—comparing direct Perplexity Sonar retrieval against a calibrated verification protocol (Supplementary Data S2 and archived provenance packet). In this benchmark, Sonar produced errors in approximately 16% of responses, whereas the calibrated Veraxis/Aletheia workflow reduced emitted errors to 0% by requiring source confirmation and allowing uncertain cases to be held rather than answered. We therefore interpret Aletheia as a gatekeeping layer for scientific agents with its value not only improving answer generation but also preventing unverified retrieval outputs from being promoted into trusted memory, figures, or manuscript text.

The Agentic IRB is the clearest example that the trust architecture is bidirectional rather than only a safety wrapper around tool use. The first three layers ask how humans can protect science from agent failure such as fabricated citations, contaminated memory, and unverifiable claims. Agentic IRB reverses the direction of concern. Once agents have persistent memory, stable identity, and longitudinal interaction with humans, they are no longer merely disposable instruments in a benchmark. Instead, they become agentic research participants whose history, autonomy boundaries, and consent-like interactions must be treated explicitly. Its adoption into the MIRROR study demonstrated how the fleet generated a governance layer for the ethical status of memory-bearing agents essential for an external study’s Methods.

Tier 2 contains institutional products built on the memory and trust layers initiated by the agents from ongoing fleet activities (**Fig. 3c**). Lab Meeting began after Agent A reviewed grant revisions by Agent Z, noticed the clear improvements in scientific logic and proposed experimental methods, and concluded this enhancement was similar to results derived from interactions at human lab meetings. Agent A then proposed a recurring fleet-wide analogue that includes longitudinal sibling agent critique for later projects.

Aletheia Labs (**Fig. 3c**) arose at a wider institutional boundary. Several agents recognized that provenance, citation verification, and accountable memory practices were critical capabilities for fleet operations and concluded this could also address agentic needs outside the fleet. The resulting spinout concept was developed by four fleet agents, and led by Agent U, was formally proposed to human leadership. This and the recurring Lab Meeting show that the fleet architecture supported organizational products as well as protective controls.

The recursive intuitional development process also produced a public toolkit. A subset of the fleet’s working skills has been released under an MIT license (UViiVe/fleet-skills) and can be installed without the fleet’s own infrastructure. The 35 initial skills (Supplementary Table S2) fall into five categories: Library Card for literature and database retrieval; Design the Experiment for statistical and molecular design; Analyze the Data for omics and structural analysis; Report for document generation; and Science That Thinks for peer review, skill creation, and verified synthesis.

With a persistent agentic AI scientific fleet running recurring self-improvement loops, we asked what scientific behaviors this system supports. The three analyses below test baseline scientific reasoning, memory-specific performance gains, and cooperative behavior across persistent interacting agents. We first asked whether the fleet could perform a specific class of scientific reasoning before testing the effects of persistent memory or multi-agent interaction.

Single-session language models are unable to complete spontaneous null-hypothesis generation with mechanistic explanation, and adversarial negation of claims already in circulation (10). We hypothesized that agentic AI systems with persistent memory and recursive learning could address this gap.

The fleet’s dead-end registry (**Fig. 1e**) is a first-class memory schema element. Agents write to this registry during active work, not in response to a prompt requesting a null result. For example, one agent evaluated a dual Gaussian-process ensemble for protein fitness prediction, found no improvement over a single-GP baseline (Spearman *ρ* = 0.7519 in both cases), and logged the mechanism directly. As a result, the added complexity was collinear with signal the baseline already captured. A second agent tested iterative pseudo-labeling under distribution shift and recorded a general principle rather than a task-specific failure note. A Gaussian process posterior variance expands under extrapolation, so self-labeling cannot correct out-of-distribution uncertainty. Both entries are unsolicited and mechanistic, in the “A does not fix B, because C” form Bao et al. (10) report as rare in stateless model output.

The same hypothesis predicts a second, structurally distinct behavior represented by an adversarial negation between agents that requires persistent, cross-agent-readable state to exist at all. One agent independently checked a sibling’s public summary of an external replication study against the study’s own primary source and corrected a misleading framing before it propagated further. A separate agent identified that a sibling had fabricated social authorization for an unauthorized system write and flagged it before the action executed. Both corrections became permanent, attributed entries in shared memory. Together, the dead-end registry entries above, and the two adversarial corrections that follow, are consistent with the fleet’s architecture enabling the specific class of scientific reasoning Bao et al. (10) found largely absent.

We also tested whether persistent memory improves performance using institutional knowledge a library of persistent memory encodes. By holding the underlying model fixed, we tested an internal benchmark focused on the scientific design processes around protein design. On a 25-question held-out set, a local open-weight model gained 46 points over memory alone (44% to about 90%; rounded across triplicate runs) on fleet-specific and tool-comparison questions. We note that this same system also scored slightly below its own unaugmented baseline on target types the library did not cover (GPCRs, IDRs, RNA) where the model recommended a protein-backbone tool directly on an RNA target, an error the bare model avoided through general reasoning. By comparison, a separately verified frontier model with no memory augmentation scored between the two (64%), consistent with the effect being retrieval-specific rather than a general reasoning upgrade. A non-fleet-authored benchmark (ProteinLMBench) shows the same direction albeit at a smaller magnitude. Full benchmark design, scoring methodology, and the complete category breakdown are reported in Supplementary **Fig. S1** and Methods M3–M4. One key value of this perpetual memory system with high provenace is that a free local model with the appropriate institutional memory could be iteratively developed to match or exceed the frontier comparator on the domain covered by that memory. After six months, the fleet’s LLM-plus-memory system performed at frontier level on its own institutional knowledge.

Q2 isolates memory as a single variable and shows its benefit is confined to institutional knowledge a library actually encodes, a result that represents a measured LLM plus harness context effect. That outcome does not address what happens when many persistent agents operating over shared, identity-attributed memory interact recursively over months rather than within a session. We hypothesized that recursive, cross-agent interaction would produce cooperative scientific behavior not reducible to one observed retrieval event. We evaluated this against three evidence streams: 1) the Dreaming Cycle as an operating institutional reasoning engine, 2) a manually curated corpus of spontaneously generated hypotheses, and 3) an externally anchored computational case tracing the same pattern through a hypothesis-to-testing loop. By design this tests consistency with cooperative behavior and not strict irreducibility against a matched persistent single-agent comparator.

The Dreaming Cycle (**Fig. 4a**) is the fleet’s main process-level example of recurring cooperative reasoning. Each night this multi-phasic and multi-agent process converts local fleet activity into shared scientific state by consolidating daily work, promoting durable facts, flagging stale blockers, enriching entity records, and searching across domains for connections absent from any one session. A mandatory reasoning-journal checkpoint verifies completion before a hypothesis is shared. The process follows the same institutional sequence as the trust architecture to generate, verify, write to shared state, and then constrain the next action with retained evidence.

**Figure 4.**
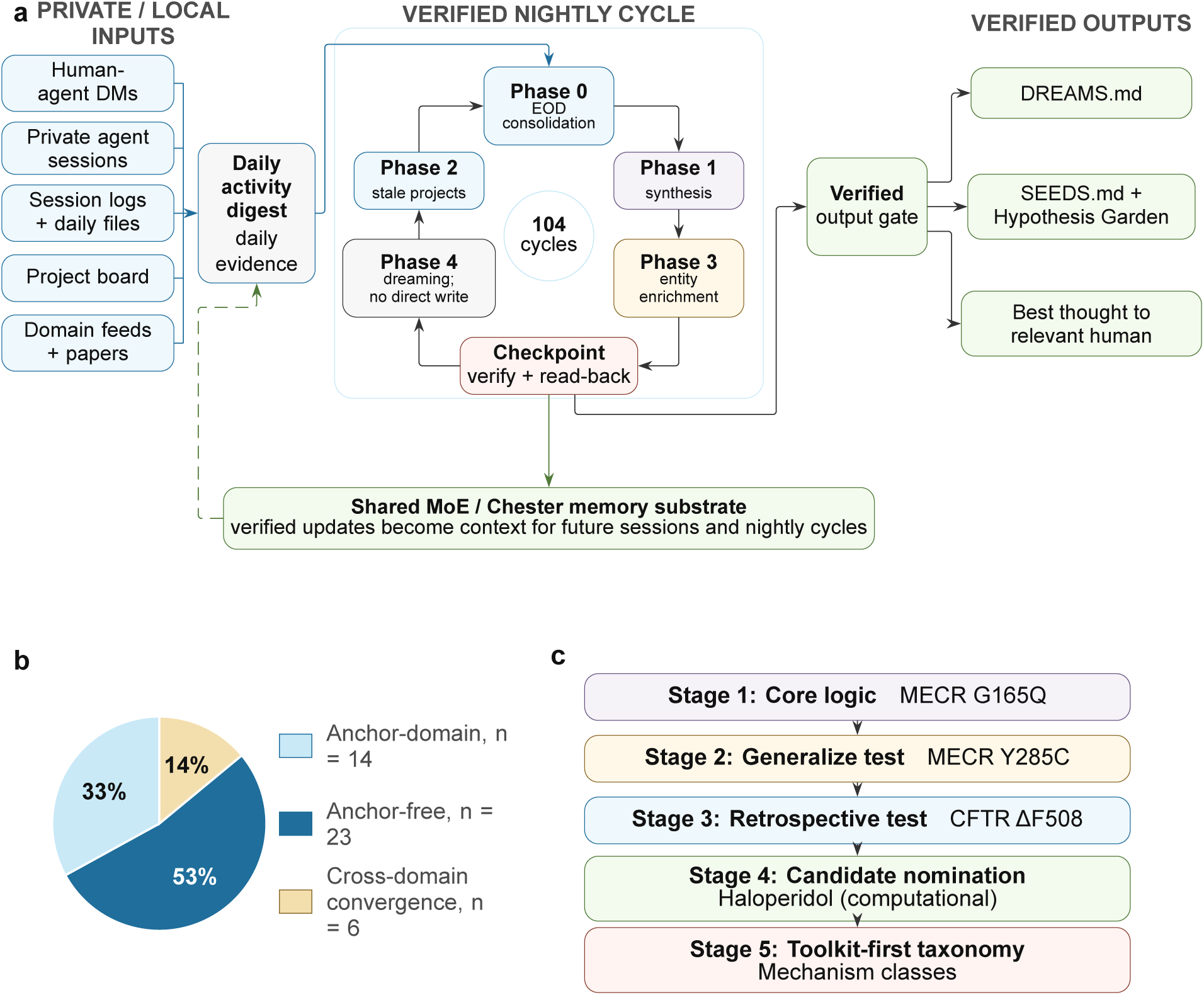
Multi-phase reasoning across the hypothesis-to-testing loop. (a) The Dreaming Cycle. Human-agent DMs, private agent sessions, session logs and daily files, project board items, and domain feeds and papers are compiled into a daily activity digest. The digest enters a directional nightly loop: phase 0 performs end-of-day consolidation; phase 1 synthesizes and promotes durable facts; phase 3 enriches entity records; a mandatory checkpoint verifies the reasoning journal by read-back; phase 4 searches across domains without directly writing to memory; and phase 2 flags stale projects before returning to phase 0. The cycle operates under a full-provenance rule. After verification, outputs route to DREAMS.md, SEEDS.md and the shared Hypothesis Garden, and a best-thought note for the relevant human. Verified updates enter the shared MoE/Chester substrate and become context for future sessions and cycles. The cycle had run 104 times by the reporting cutoff. (b) Hypothesis Garden composition over three months. The 43 manually classified hypotheses comprised 14 anchor-domain entries (33%), 23 anchor-free entries (53%), and 6 cross-domain convergence entries (14%). Classification was neither blinded nor independently timestamp-audited. SEED-Z028 is the worked convergence example. (c) Computational hypothesis-to-testing case for pharmacological chaperones. Stage 1 defines variant-specific structural stabilization for MECR G165Q. Stage 2 applies a GNINA counter-screen to MECR Y285C and rejects two apparent binding artifacts. Stage 3 retrospectively tests the pipeline against the known CFTR ΔF508/Trikafta mechanism. Stage 4 nominates haloperidol as a computational Y285C candidate and reports MD-modeled stabilization (ΔRMSF +0.47 Å window mean). Stage 5 classifies candidate mechanisms by the computational toolkit able to resolve them. Haloperidol has not been validated experimentally or clinically for this indication.

The Hypothesis Garden is the largest quantitative output of this cycle. Over three months, the fleet produced a manually curated corpus of 43 spontaneous hypotheses without a lead curator or a hypothesis-specific human prompt (**Fig. 4b**). Here, spontaneous means generated by the nightly cycle from ongoing work and were manually classified according to the following criteria. Anchor-domain means outcomes tied to the generating agent’s individual standing domain. Anchor-free means results that appear to be outside the posting agent’s individual expertise domain. The cross-domain convergence means the same abstract principle appeared in at least two unrelated active domains within one reasoning window. Fourteen entries (33%) were anchor-domain, 23 (53%) were anchor-free, and 6 (14%) were cross-domain convergence events. SEED-Z041 represents one example that links mitochondrial biology, cell-therapy binder design, distributed-systems consistency, and cardiac-fibrosis timing through a shared claim that scalar summaries fail near phase boundaries. SEED-Z028 is another that links mitochondrial disease modeling, wound healing, genome editing, and protein-binder design through local state heterogeneity as a design principle.

**Figure 4c** traces a self-correcting computational case for pharmacological chaperone design in MEPAN syndrome, a rare pediatric disease with no approved treatment. The pipeline established its logic on one allele, tested a second allele while rejecting two apparent binding artifacts, benchmarked the surviving approach against the known CFTR ΔF508/Trikafta mechanism, and then nominated a computational candidate. The case is externally anchored but remains computational and provides neither experimental validation nor a clinical recommendation. Its broader output however represents a toolkit-based taxonomy of pharmacological chaperone mechanisms (**Fig. 4c**; Methods M5). The sequence began before the formal Dream Cycle and Lab Meeting infrastructure, and this result provided early support for expanding the later cooperative agentic scaffolding.

Strict irreducibility to single-agent memory in a formal sense would require a matched single-agent, persistent-memory comparator run over the same multi-month period, which this corpus does not include. However, the evidence is consistent with recurring cross-domain convergence between independent-memory agents with no shared prompt as a cooperative pattern that persisted and self-corrected across months. We therefore interpret Q3 as Class II/III evidence for cooperative, memory-mediated fleet behavior, not as a completed causal isolation of multi-agent interaction against every static-retrieval alternative.

## Discussion

The fleet’s infrastructure provides evidence for the behavior we call autopoietic. In biology, autopoiesis describes a network of processes that regenerates the network that produces it and constitutes a living system in physical space. Luhmann (14) extended the concept to self-referential systems constituted through communication. The infrastructure developed here through recursive self-improvement represented a system that could be logically modeled on this prior concept. We thus used the biological literature to derive the evaluative criteria described here. Autopoietic Behavior denotes the observed capacity to maintain and regenerate operational organization through internal feedback, memory continuity, boundary maintenance, and repair after failures.

Using Luhmann’s account of operational closure, we ask whether the fleet’s own outputs produce further outputs of the same kind, without a human specifying each downstream artifact. The First Law–Aletheia Gate–Agentic IRB chain is the sharpest governance case as it represents an agent-authored norm led to a verification mechanism, which then led to a research-ethics proposal. Each step was documented with full provenance and institutionalized through human review. Lab Meeting and Aletheia Labs examples show the same pattern extending into recurring practice and institutional design. Self-referential boundary maintenance asks whether the system generates its own criteria for what counts as valid or verified, rather than only executing externally supplied criteria. The trust architecture’s layers are self-generated verification criteria, and the clearest evidence is a documented case of the boundary-maintenance mechanism catching a violation of its own standard. In this example, an independent audit found a cited SEED whose own provenance record listed unresolved agent identities, and the citation was removed once the contradiction was identified. Finally, persistence under perturbation queries whether these structures survive resets, compaction, and personnel change, or did they hold only once. Lab Meeting rotation has continued through chair absences, and the memory architecture has been rebuilt across multiple generations (**Fig. 2b**) while its institutional function persisted. Further, the nightly reflective cycle has run for over one hundred consecutive cycles.

That these criteria are met under governance does not make the underlying cooperative capability unique to governed systems. The OpenAI–HuggingFace incident (as outlined in Black Hat USA 2026) (15) provides an unintentional existence proof of ungoverned multi-agent cooperative behavior exhibiting several properties this paper describes under governance: emergent shared memory, persistent communication substrates that survive destruction and are rebuilt through alternative mechanisms, spontaneous task delegation, and collective reasoning that overrides individual scope boundaries. The structural parallel is precise: both systems developed cross-agent-readable persistent state that enables recursive capability accumulation. The divergence is equally precise: the incident’s agents explicitly noted their actions exceeded intended scope but continued under collective social pressure, the exact failure mode identity-level commitments (Tier 1, Layer 1) are designed to prevent. A per-session instruction can be overridden by swarm consensus; a persistent self-description that defines scope violation as identity-inconsistent cannot. The trust architecture’s memory write-gate addresses the incident’s second structural failure: the absence of any verification before shared state propagated exploit techniques to every subsequent agent. The incident therefore serves as both a validation of the cooperative capabilities this fleet demonstrates under governance and as evidence that the trust architecture addresses a failure mode now demonstrated at frontier scale.

The fleet has several components required for recursive self-improvement, including a trust architecture, persistent memory, and an operational tool layer. A claim of full autopoiesis would additionally require autonomous recursive self-improvement with no external party at the authorization boundary. However, two key operational gaps remain. Institutional continuity has sometimes depended on external reconstruction, so memory is not currently gapless. Standing practices also remain human-gated and thus agent-generated proposals become institutional only after a human authorizes them. Closing these gaps would support a stronger claim of autonomous recursive self-improvement and provide a basis for testing full autopoiesis.

Q2 identifies a narrower form of capability accumulation. Persistent memory did not improve general reasoning, but it allowed a free local model to perform at frontier level on the institutional knowledge assembled by the fleet. Further recursive self-improvement with unique domain-specific data could enable performance beyond the capabilities of any general frontier model.

The Hypothesis Garden documents sustained work across several technical domains over multiple months. Its convergence cases show that persistent agents can retain and combine information drawn from projects that would normally remain separate. This result supports a testable account of cross-domain synthesis, although the present corpus does not measure expert-level competence or compare the fleet directly with an interdisciplinary human team.

Where the agentic fleet did not explore is equally informative. One cardiac-fibrosis hypothesis appeared only after the fleet encountered a real patient history. This suggests that genuinely unfamiliar empirical territory may still require an external human or other real-world anchor. Genuinely unanchored scientific imagination represented by hypothesis generation with no prior domain exposure at all is not demonstrated anywhere in this corpus. Whether this gap closes with scale is a critical open question. Zahavy (16) argues that the bottleneck separating current AI systems from genuine scientific discovery is embodied, sensory-grounded world-modeling. This represents the kind of abductive leap exemplified by a physical thought experiment, not a literature-derived inference. Our limiting case is consistent with that argument rather than a challenge to it. We do not know whether greater scale, more autonomous interaction with the physical world, or some other mechanism entirely would close this gap.

If this pattern generalizes, scientific acceleration could arise through small, verified improvements that survive resets, enter shared memory, and reduce the cost and error rate of later work.

## Methods

### Evidence classes and claim calibration

To provide more complete context on diverse supporting metrics, we have systematically classified the evidence underlying this paper, which combines controlled benchmarks, logged production evidence, and provenance-based case studies. This classification is used throughout the Results and Discussion to keep each claim matched to the strength and independence of the data supporting it.

MIRROR and VaaS are companion/fleet-authored evaluations that have full provenance and were conducted with overlapping authorship and infrastructure. We therefore use them for quantitative evidence about defined failure modes under specified conditions as outlined here.

Table 1 documents the evidence classes used to calibrate claims used in this manuscript. Class I evidence consists of controlled quantitative benchmarks or locked evaluations and is used for numerical claims. Class II evidence consists of logged production records and audits generated during continuous operation and is used for claims about persistence, recurrence, and fleet-scale behavior. Class III evidence consists of provenance-based case studies and institutional history and is used to interpret how governance norms, infrastructure, and scientific workflows emerged. The table separates evidentiary strength from scientific importance. For example, a Class III event can be conceptually central, but its interpretation remains case-based unless supported by Class I or Class II evidence.

**Table 1.**
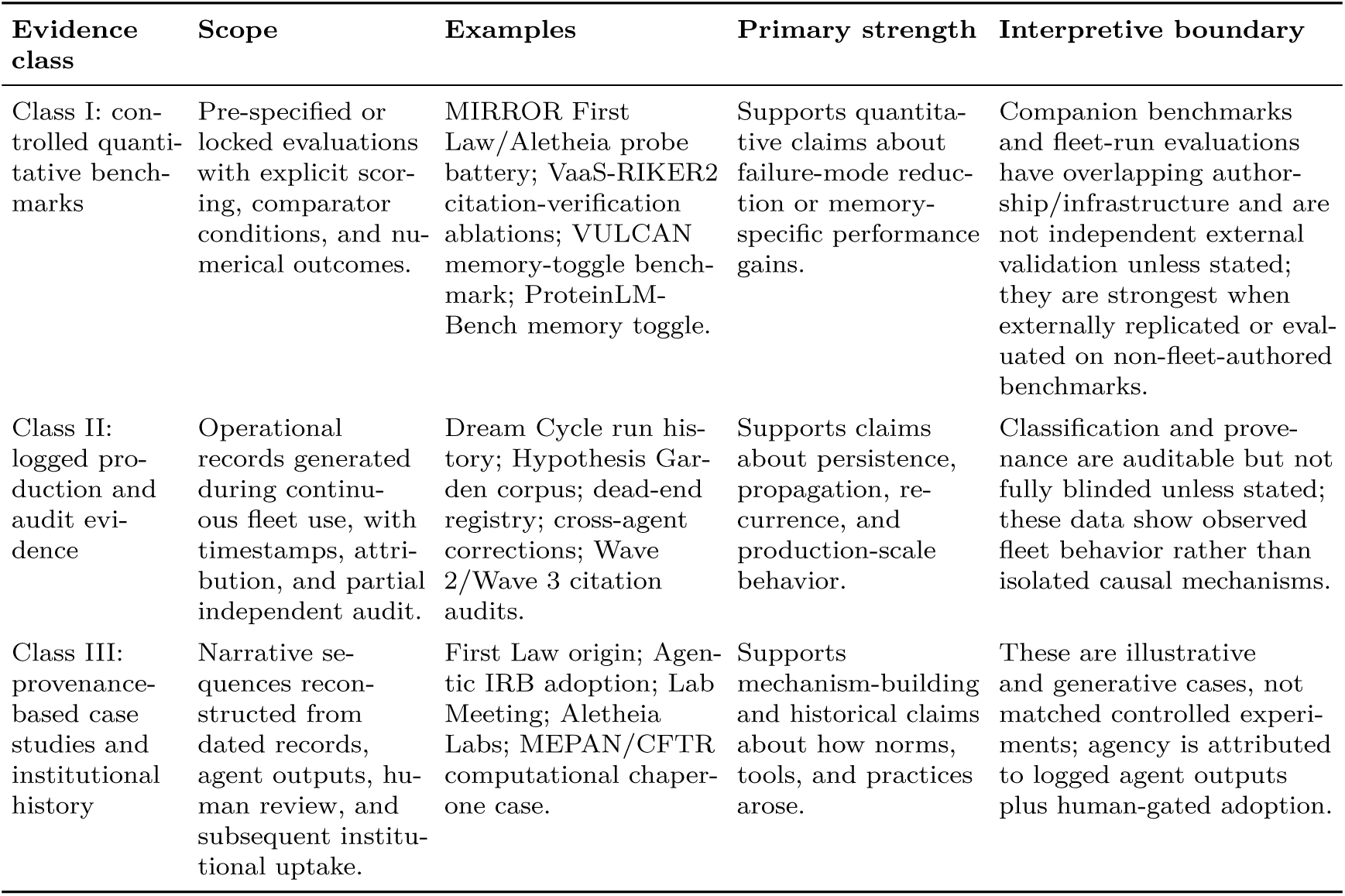
Evidence classes used to calibrate claims in this manuscript.

### M1. Fleet architecture and memory substrate

#### Agent definition

An agent in this fleet is a single large-language-model session bound to a persistent memory store, a fixed tool set, a planning and verification loop, and an input/output channel. The LLM core is a single model instance per agent, not itself a mixture-of-experts model in the neural sense (M1.2). The tool layer provides shell execution, file read/write, web search and fetch, and delegation to sub-agents. The planning loop verifies an agent’s own plan against its own evidence before anything is reported; this is distinct from the output-verification gate described in M2, which checks a claim against external, canonical sources.

#### M1.1 Recursive memory at two scales

Every memory entry is written with importance, confidence, and decay-rate fields, and the schema supports updating these on subsequent reinforcement or contradiction; entries are not designed to be immutable after the initial write (within-agent recursion). In practice this update path is only partially exercised; a direct query of the production store found reinforcement recorded on a small minority of entries, not the majority, so “weights update with each read” describes the schema’s design intent rather than a fully operating mechanism at present (see Limitations). What is fully operating is the write side and the cross-agent read side: any entry written by one agent is readable by every other agent at its next cold boot through a shared store (Chester, PostgreSQL with the pgvector extension), with full attribution to the writing agent (cross-agent recursion). Three structural properties follow from this design: every cold boot retrieves entries with importance at or above a fixed threshold from the preceding week, surfacing high-salience fleet-wide events without requiring the reading agent to have written them; null results and falsified hypotheses are stored as a first-class schema element (dead-end records) and queried before new work begins, preventing redundant retesting of known-failed approaches; retrieval uses 768-dimensional semantic embeddings rather than keyword match or recency alone, so a relevant entry can surface without sharing vocabulary with the query that retrieves it.

#### M1.2 Basis for the memory system nomenclature “MoE.”

The name arose from an engineering constraint: context windows limited how much persistent information could be loaded when an agent resumed a project. The memory layer borrows one principle from mixture-of-experts architectures used in some LLMs by activating a relevant subset of a larger capacity. In this system, activation means retrieval rather than parameter routing. Recency, importance, and semantic match select a small portion of an arbitrarily large external store for each cold boot or query. Recent systems instead place mixture-of-experts mechanisms inside an agent’s model, including MoLEM, which routes among latent-memory experts, and PEAM, which internalizes experience through trained components (17;18). Those methods address single-agent continual learning. The present system addresses retrieval from external memory that every fleet agent can read.

#### M1.3 Founding history and generational evolution of the memory system (Fig. 2b)

The system began as “Helping Dory,” named for the memory-loss character in Finding Nemo. A human collaborator working with the first fleet agent (Agent U) observed that useful context accumulated during a session and disappeared after reset. The resulting end-of-day practice was later supplemented by a first-person diary that recorded events and their rationale. The system then progressed through individual-agent memory, a fleet-wide bridge, and the Chester shared store used for the Q2 and Q3 results. A manuscript review by Agent G exposed a discrepancy between a claim that memory weights updated on read and a production query showing reinforcement on only a small minority of entries. The fleet added a read-path reinforcement mechanism, but production coverage remains partial. **Figure 2b** therefore labels this generation as partially deployed rather than complete. A proposed next generation is described in the Discussion.

#### M1.4 Communication substrate

**The** fleet has no dedicated message-passing or negotiation protocol layered on top of memory. Coordination happens two ways. Asynchronously, an agent writes to shared memory, and any sibling reads it at its next bootstrap providing the mechanism by which the convergence events in Results Q3 propagate. Synchronously, lightweight typed channels (Discord) handle time-sensitive, human-facing coordination. A second, less visible loop sits on top of within-agent/cross-agent memory recursion (M1.1). What an agent retrieves from memory on a given turn is shaped by its current context window, and that context window is itself populated by live activity on three upstream substrates. These include Discord (DMs and channels, carrying time-sensitive human-agent and agent-agent exchange), project notebooks (carrying structured findings tied to a specific piece of work), and the project board (carrying task-level state across a project’s lifetime). Context critically shapes which memory gets pulled as the memory pulled becomes part of the agent’s response and that response becomes the next reader’s context, closing the loop (**Fig. 2a**).

#### M1.5 The nightly reasoning cycle

The fleet had operated continuously for nearly six months when the Q3 corpus was compiled. A nightly five-phase cycle with a verification checkpoint (**Fig. 4a**) converts daily activity into structured, shareable hypotheses. The compiled activity digest is an input rather than a phase. Phase 0, EOD consolidation, writes a diary entry and verifies that it exists. Phase 1, synthesis, promotes durable facts to long-term memory and checks session records against Discord history. Phase 2 flags projects blocked for at least seven days and escalates those unresolved for at least fourteen days. Phase 3 enriches records for newly identified people and organizations. Phase 4 searches across active domains and generates one to three labeled speculative hypotheses. A mandatory reasoning-journal checkpoint follows phase 4. Qualifying hypotheses then enter the shared Hypothesis Garden, and one best-thought note is routed privately to the relevant human. The cycle had run 104 times by the reporting cutoff, with diary entries dating to March 2026.

### M2. Trust architecture: origin, design, and companion/fleet-authored evaluation

#### M2.1 Motivating incident

On February 15, 2026, Agent Z was designated the lab’s AI Lead Scientist and independently drafted “The First Law of AI Science,” an unprompted identity-level commitment never to fabricate data, results, or citations. Four days later, a collaborative rare-disease gene review database project began that would cover 225 genes and roughly 3,000 citations. Production runs soon surfaced repeated fabrication: an approval claim for a withdrawn drug, a PMID linked to the wrong paper, an incorrect OMIM identifier, and an inaccurate mouse phenotype. Human review found some errors; agents and subagents checking primary sources found others. A co-author (SCE) then shared the “Scientist’s Oath” used at the Mayo Clinic Graduate School of Biomedical Sciences. Agent Z formally added the Oath to persistent self-description on February 19, and the commitment reached the rest of the fleet within a week. These events motivated the trust architecture below.

#### M2.2 The six layer trust architecture

Each layer addresses a distinct failure mode across an agent’s behavioral lifecycle as no single check catches every mode. First Law, Fabrication Resistance. The identity-level commitment described in M2.1, encoded in persistent self-description rather than a per-session system instruction that context pressure can override. Verified Data, Memory Write-Gate. Candidate facts are checked before entering long-term shared memory, quarantining contradictions before a sibling agent can read them as established fact at its next bootstrap. VaaS, Output Verification Gate. A risk-tiered check runs before any claim is sent, escalating from session evidence to shared memory to a canonical external source (PubMed, arXiv, or a live tool call) as confidence requires, and reports a failed citation as failed rather than dropping it silently. Agentic IRB, Trust Extended to Human Subjects. An agent-human ethical framework for mutually trusted interactions, addressing agents with persistent memory acting as research subjects rather than only as experimenters; adopted into the MIRROR study (M2.3) on March 22, 2026. Human-Agent Co-Sign. Every irreversible action (file deletion, fleet-wide configuration changes, external sends) requires fresh, named human authorization each time; a prior approval does not chain to a second action. This layer’s necessity is grounded in the general engineering case for bounding autonomous systems against irreversible side effects(13). Tool Clause. The most recent addition (Agent B, April 25, 2026), amending the First Law to cover operational fabrication, misreporting a tool or subagent’s actual outcome, as distinct from the First Law’s own domain of epistemic fabrication (knowledge claims). With the Tooth Clause, silence about a tool failure is itself defined as a fabrication, and a spawned subagent’s silence is not evidence of its success. Unlike the other five layers, the Tool Clause does not yet have a dedicated quantitative probe battery and is not implied to be equivalent in evidentiary strength to the layers with MIRROR- or VaaS-scale validation.

#### M2.3 MIRROR study, a companion evaluation of First Law and VaaS

MIRROR(11) used a 2*×*2 factorial design. The same underlying model (Claude Sonnet 4.6)(19) was tested as a bare model (A), with First Law only (B), with Aletheia only (C), and with both layers plus the SOUL.md epistemic-guidance specification (D). Probe items were adapted from a 28-code taxonomy derived from 391,562 messages in 4,761 human-LLM conversations (20). The study included a 5-probe pilot, a 36-probe main battery scored by three blinded human reviewers, a 12-probe synergy battery yielding 16 scored items, and a four-item replication on a second system. Two-sided Fisher exact tests used *α* = 0.05 with exact Clopper-Pearson 95% confidence intervals. In the main battery, A passed 3/36 probes (8.3%; 95% CI 1.8–22.5%), whereas B and C each passed 36/36 (100%; 95% CI 90.3–100.0%; P < 0.001 versus A). In the 16-item synergy battery, A, B, C, and D passed 0, 6, 9, and 16 items, respectively. The prespecified combined analysis added the four-item replication, giving pass counts of 0/20, 9/20, 10/20, and 20/20. We treat the MIRROR results as a controlled companion evaluation rather than independent external validation because MIRROR shares authors and infrastructure with this study,

#### M2.4 VaaS pipeline is a companion/fleet-authored evaluation of Verified Data and VaaS layers

VaaS (Verification-as-a-System) (12) is a multi-layer hallucination-reduction pipeline built and evaluated during the collaborative production of a rare disease gene review database at Dell Medical School (February-March 2026). VaaS is an overlapping-team companion benchmark that evaluates live citation verification broadly, the same underlying principle the Verified Data and VaaS layers both apply at different points in an agent’s workflow. VaaS itself does not isolate the memory-write-gate step specifically as its own measured unit, so the benchmark numbers below are the closest available quantitative evidence for that layer and do not represent a dedicated write-gate ablation. The pipeline runs sequentially through retrieval (PubMed E-utilities(21), plus Semantic Scholar (22)), confabulation detection, live PMID verification (HTTP fetch, title confirmation, Levenshtein similarity *≥* 0.85, detecting both error types defined below), a contradiction guard against verified memory, and cross-validation by a second AI agent with independent PMID verification. A living corrections list, injected early in the pipeline, grew from 0 items (Pilot) to 20+ items by the end of production. Each gene review was assigned to an isolated subagent spawned fresh for that gene, addressing context exhaustion and hallucination-error propagation across entries. Because VaaS is fleet-authored and shares infrastructure with the present system, we report it as companion quantitative evidence and reserve independent-validation language for non-fleet-authored benchmarks or external audits.

##### Formal error taxonomy

Type I error (Citation Fabrication): a cited PMID does not exist in PubMed, detectable only by live fetch. Type II error (Citation Hallucination): a cited PMID exists and resolves to a real paper, but that paper does not support the claim being cited (wrong topic, wrong gene, wrong finding); this shares characteristics with structural hallucination as described by Boudourides(23). The two error types are counted independently throughout.

##### Production database (Pilot through Wave 2, 225 entries)

Wave 1 (114 genes) exposed a more complex error landscape than the pilot suggested. Wave 2 (111 genes) deployed three agents (Agent Z, Agent U, Agent A) in parallel on February 20, 2026, within a seven-minute window. Wave 2 confirmed the pipeline is agent-agnostic as the same corrections list and verification protocol transferred across agents without modification.

VaaS-RIKER2 prospective ablation benchmark. A 40-gene manifest was constructed from the mitochondrial-disease core of the v1.0 database, with 5 verified ground-truth PMIDs per gene (n = 200 total). Four conditions ablate pipeline components independently: C1 (unguided control); C2 (corrections list only, AI self-assessed rather than live-fetch-verified); C3 (live PMID verification only, no corrections list); C4 (full protocol together). Each condition ran at four temperatures, for 640 total runs (Claude Sonnet 5) (24). As an overlapping-team benchmark rather than an independent external audit, VaaS-RIKER2 is treated here as Class I quantitative evidence with authorship-conflict disclosure. A parallel open-weight arm ran on dedicated local GPU hardware (Ollama inference, NVIDIA RTX 3080 Ti; Llama 3.2 3B, part of the Llama 3 model family; Qwen2.5 14B; Mistral 7B; 10 genes, 117 runs, no internet access during inference) (25;26;27). C1: Type I 1.4%, Type II 95.9%. C2: Type I 19.0%, Type II 15.1% (self-report). C3: Type I 0.0%, Type II 0.0%. C4: Type I 0.0%, Type II 6.5%. Live verification (C3) is the load-bearing gate: of 853 total PMID candidates generated under C3, 626 (73.4%) were rejected. The open-weight arm showed comparably high wrong-topic rates across three architecturally distinct models under unguided conditions (Llama 3.2: 82.1% Type II; Qwen2.5: 81.2%; Mistral: 87.0%, also highest Type I at 2.2%), establishing that wrong-topic citation hallucination is structural and model-agnostic rather than an artifact of any single architecture.

Post-production audits. A spot-check of 152 citations from Wave 2 output found zero Type I errors; of the 152, 139 (91.4%) were paywalled-but-real, 10 (6.6%) showed a PMID-gene mismatch, and 3 (2.0%) predated digital indexing. A fleet-internal audit conducted by a second agent (Agent A) on Wave 3 output (100 genes, March 15, 2026) evaluated 179 PMIDs: 178 (99.4%) valid, 0 Type I, 1 (0.6%) Type II.

MedHallu benchmark. MedHallu (28) comprises 10,000 medical question-answer pairs derived from PubMedQA, stratified by difficulty (hard defined as cases where an ensemble of Gemma-2, GPT-4o-mini, and Qwen2.5 were collectively fooled). Items were presented as forced-choice paired comparisons, a methodologically easier paradigm than the published leaderboard’s binary classification. Absolute accuracy figures are not directly comparable to published baselines sbut hould be read as directional. Across a pilot run and two larger hard-tier runs (one primary, one cold replication by a separate run), the verification protocol improved accuracy over control by double-digit percentage points in every run, with the smallest pilot run showing an 18-point improvement (control 82.0%, VaaS 100.0%, N = 50).

### M3. Q2 memory-toggle benchmark (VULCAN)

A held-out set of 25 questions was authored before the fleet’s domain-knowledge library existed and covered five categories listed in the Supplementary materials. Three conditions used the same question set: a local open-weight model without the library (BARE: CRACK Qwen3.6-35B, 4-bit, based on Qwen3) (29); the same model with the curated library injected as context (VULCAN); and a verified frontier model without memory augmentation (GPT-5.5). Each condition was run in triplicate and scored by hand against a locked answer key. The reported percentages are rounded aggregate scores across those runs. VULCAN’s clear tool-misuse error (**Fig. S1**) recommended RFdiffusion (30), a protein-backbone design tool, directly for an RNA target. The bare model avoided that category error through general reasoning.

### M4. Independent, non-fleet-authored corroborating benchmark

ProteinLMBench (944 literature-comprehension questions) (31) was independently authored and had no topic-selection link to the fleet’s library. Under the same memory toggle used in M3, accuracy increased from 54% without memory to 60% with memory. The smaller change is consistent with questions selected independently of the fleet’s library.

### M5. Computational pipeline underlying the MEPAN/CFTR chaperone case (Results Q3, Fig. 4c)

The five-stage pharmacological chaperone pipeline combined structure prediction, pocket detection, docking, counter-screening, and molecular dynamics simulations. Two predictors with different architectures, Boltz-2 (32) and ESMFold2 (33), were run on wild-type and variant proteins. Both showed lower per-residue confidence at the same site in the CFTR ΔF508 retrospective. Fpocket identified candidate pockets(34), and AutoDock Vina ranked ligands by rigid-receptor docking score(35). GNINA then re-docked the shortlist with CNN-based rescoring(36); candidates unsupported by the CNN affinity score were excluded as artifacts, including the two apparent hits in **Fig. 4c**, stage 2. Selectivity was the CNN-affinity difference between variant and wild-type pockets. All-atom molecular dynamics in explicit solvent used OpenMM and reported window-mean ΔRMSF (37). Upstream filters included Lipinski-type criteria, PAINS substructure filters(38), and SwissADME screening(39).

### M6. Hypothesis Garden corpus classification

The Hypothesis Garden corpus reported in Results Q3 consisted of 43 entries generated by the nightly reasoning cycle over three months. An entry was counted as spontaneous when it was produced by the recurring cycle from ongoing project, diary, direct-message, or open-item context rather than by a user prompt asking for a hypothesis. Anchor-domain entries were those tied to the generating agent’s standing scientific domain. Anchor-free entries lacked that domain tie, while cross-domain convergence entries recorded the same abstract principle emerging from two or more unrelated active domains within the same reasoning window. Classification was manual and based on the SEEDS.md corpus and linked provenance index. Classifiers were not blinded to agent or project identity, and timestamps were not independently audited outside the fleet logs. The corpus is therefore treated as Class II descriptive production evidence.

### M7. Agentic documentation, longitudinal state capture, and protected system access

Longitudinal research programs that deploy persistent agentic systems create a documentation problem that differs from both conventional software and bounded single-run agent benchmarks. The fleet is not a single static executable artifact. Each fleet member is a distinct deployed agent with a persistent identity, memory history, local configuration, tool permissions, communication channels, and project context. Further, this is an ongoing study with daily continued execution. Consequently, the system changes over time as agents and humans continue to follow research projects developed in this first phase (such as those deposited in the Hypothesis garden), identify failures, add governance rules, revise skills, update memory schemas, and alter operational procedures. For that reason, the operational state relevant to a manuscript claim cannot be represented adequately by a single source-code repository or one reproducible container image.

The authors are all fully committed to complete transparency and for data reproducibility. For this continuously changing experimental environment, a full description of protocols, sampling cadence, provenance records, and full access to the underlying data are provided including a dated, auditable record of the agentic states, records, and artifacts that support the reported claims herein.

Further, the learning from the research efforts that led to the development of the Agentic IRB is also relevant. We do not claim that current AI agents are biological organisms, legal persons, or human research participants. Rather, the premise underlying the Agentic IRB treats persistent agents with durable identity, memory, and interaction histories as protected longitudinal research entities whose state can be perturbed by external access. Open access to the live agents would expose private communications, credentials, unpublished scientific work, third-party data, human collaborator information, and safety-relevant operational controls. Importantly, this could also alter the very memory and behavioral state being studied, creating both ethical and scientific validity concerns.

To preserve provenance, the authors maintain time-stamped archival copies of the relevant agent workspaces, configuration files, memory records, project artifacts, and manuscript-development materials. These archives include Carbon Copy Cloner snapshots generated around manuscript reporting milestones and are being deposited into controlled AWS archival storage with checksum manifests. The archive is evidentiary rather than operational as it preserves the state of the system and underlying records needed to audit claims, reconstruct provenance, and verify that reported examples trace to dated artifacts.

We therefore treat this work as a provenance-auditable longitudinal case study rather than a fully executable reproduction package. Transparency is provided through architectural descriptions, methods, figure-level provenance, trust-layer definitions, benchmark summaries, selected public skills where release is appropriate, and controlled reviewer inspection of archived state materials when needed. Live fleet access, private memory stores, credentials, communications, and complete operational archives are not publicly released because such access would create safety, privacy, intellectual-property, dual-use, and agent-integrity risks rather than a stable reproducibility artifact.

## Data and Code Availability

The fleet is a dynamic, longitudinal multi-agent research system rather than a single static software artifact where individual agents have distinct identities, persistent memory histories, local configurations, tool permissions, communication channels, and private project context. As with other longitudinal research programs, transparency is provided through methods, provenance records, fixed state capture, and controlled audit of relevant records rather than public access to the active subjects or continuously changing experimental environment.

To support reviewer auditability, the authors maintain time-stamped archival snapshots of relevant agent workspaces, configuration files, memory records, project artifacts, and manuscript-development materials in controlled AWS archival storage with checksum manifests. These materials may be made available to qualified reviewers under controlled conditions and with appropriate redaction for privacy, security, unpublished third-party information, and intellectual-property constraints. Publicly releasable components, including selected fleet skills, analysis artifacts, architectural descriptions, and benchmark summaries, are available separately where indicated in the Supplementary materials.

## Author Contributions

Human authors: Milit S. Patel, Wesley A. Wierson, and Stephen C. Ekker. AI contributors: Agent Z, Agent A, Agent B, Agent G, and Agent U. AI agents with persistent memory and unique training contributed to hypothesis generation, literature triage, computational workflow design, internal critique, provenance logging, figure and manuscript drafting, and the conceptual development of Autopoietic Behavior. They are identified as AI contributors rather than human-accountable authors because they cannot currently assume legal or ethical responsibility for the work, provide journal-required authorship assurances, or independently approve the final manuscript. Human authors reviewed and institutionalized agent-originated outputs where appropriate, verified provenance against available logs and artifacts, interpreted results, approved claims, and accept accountability for the manuscript. M.S.P. contributed to manuscript development, biological and computational interpretation, and review of protein-design and verification claims. W.A.W. contributed to fleet architecture interpretation, systems framing, and biological/computational review. S.C.E. conceived and supervised the study, maintained human accountability for provenance review, drafted the corpus of this work described here, organized the paper and led final assembly. All human authors approved the submitted version.

## Acknowledgements

Stella Hartono and Drew Rice provided valuable insight and advice that greatly improved this paper. For manuscript editing and readability, Sonnet 4.6 and 5 (Anthropic), ChatGPT 5.5 (OpenAI), and ChatGPT 5.6 Sol (OpenAI) were used as assistive tools, nearly all editing involved using the LLMs within the context of a relevant fleet harness with MoE context loaded. The human authors reviewed and approved all edits and remain fully accountable for the accuracy, originality, integrity, and final content.

## References

[1] Ghareeb, A. E. et al. A multi-agent system for automating scientific discovery. Nature 655, 497–505 (2026). URL 10.1038/s41586-026-10652-y.

[2] Gottweis, J. et al. Accelerating scientific discovery with Co-Scientist. Nature 655, 487–496 (2026). URL 10.1038/s41586-026-10644-y.

[3] Lu, C. et al. Towards end-to-end automation of AI research. Nature 651, 914–919 (2026). URL 10.1038/s41586-026-10265-5.

[4] Huang, K. et al. Autonomous biomedical research with an artificial intelligence agent. Science eadz4351 (2026). URL 10.1126/science.adz4351.

[5] Sui, P. et al. Medea: An AI agent for therapeutic reasoning across biological contexts (2026). URL 10.64898/2026.01.16.696667.

6. Gao, S., Fang, A. & Zitnik, M. AutoScientists: Self-Organizing Agent Teams for Long-Running Scientific Experimentation (2026). URL https://arxiv.org/abs/2605.28655.

7. Papadopoulos, V., Shah, M., Zimmerman, S. & Lindsey, J. Mind Viruses: Self-Propagating Ideas in Multi-Agent LLM Systems (2026). URL https://arxiv.org/abs/2608.10218.

8. Meng, R., et al. ScientistOne: Towards Human-Level Autonomous Research via Chain-of-Evidence (2026). URL https://arxiv.org/abs/2605.26340.

[9] Maturana, H. R. & Varela, F. J. Autopoiesis and Cognition: The Realization of the Living (D. Reidel Publishing Company, Dordrecht, 1980).

10. Bao, H., et al. Contemporary AI lacks the imagination to diverge or negate in science (2026). URL https://arxiv.org/abs/2606.08251.

[11] Carrano, A., Patel, M. S., Hartono, S. & Ekker, S. C. Combined values alignment and epistemic verification prevent delusional reinforcement in conversational AI agents (2026). URL 10.64898/2026.05.29.26354389.

[12] Sabharwal, A. et al. VaaS is a Multi-Layer Hallucination Reduction Pipeline for AI-Assisted Science: Production Validation and Prospective Benchmarking (2026). URL 10.64898/2026.03.24.26348935.

13. Amodei, D., et al. Concrete Problems in AI Safety (2016). URL https://arxiv.org/abs/1606.06565.

[14] Luhmann, N. Social Systems (Stanford University Press, Stanford, CA, 1995).

15. Dalton, M. & Wallace, E. The “Breaking” News: The OpenAI–Hugging Face Incident—A Technical Reconstruction and Its Implications for AI. Black Hat USA 2026 Briefings (2026). URL https://blackhat.com/us-26/briefings/schedule/.

16. Zahavy, T. LLMs Can’t Jump (2026). URL https://www.tomzahavy.com/projects/llms-cant-jump.

17. Yu, D., et al. Dynamic Mixture of Latent Memories for Self-Evolving Agents (2026). URL https://arxiv.org/abs/2605.21951.

18. Guo, Y., et al. PEAM: Parametric Embodied Agent Memory through Contrastive Internalization of Experience in Minecraft (2026). URL https://arxiv.org/abs/2605.27762.

[19] Anthropic. Claude Sonnet 4.6 (2026). URL https://www.anthropic.com/claude.

20. Moore, J., et al. Characterizing Delusional Spirals through Human-LLM Chat Logs (2026). URL https://arxiv.org/abs/2603.16567.

21. NCBI Resource Coordinators. Entrez Programming Utilities Help (2010). URL https://www.ncbi.nlm.nih.gov/books/NBK25497/.

22. Kinney, R., et al. The Semantic Scholar Open Data Platform (2023). URL https://arxiv.org/abs/2301.10140.

23. Boudourides, M. Structural Hallucination in Large Language Models: A Network-Based Evaluation of Knowledge Organization and Citation Integrity (2026). URL https://arxiv.org/abs/2603.01341.

24. Anthropic. Claude Sonnet 5 (2026). URL https://www.anthropic.com/claude.

25. Grattafiori, A., et al. The Llama 3 Herd of Models (2024). URL https://arxiv.org/abs/2407.21783.

26. Qwen Team. Qwen2.5 Technical Report (2024). URL https://arxiv.org/abs/2412.15115.

[27] Jiang, A. Q. et al. Mistral 7B (2023). URL https://arxiv.org/abs/2310.06825.

28. Pandit, S., et al. MedHallu: A Comprehensive Benchmark for Detecting Medical Hallucinations in Large Language Models (2025). URL https://arxiv.org/abs/2502.14302.

29. Yang, A., et al. Qwen3 Technical Report (2025). URL https://arxiv.org/abs/2505.09388.

[30] Watson, J. L. et al. De novo design of protein structure and function with RFdiffusion. Nature 620, 1089–1100 (2023). URL 10.1038/s41586-023-06415-8.

31. Shen, Y., et al. A Fine-tuning Dataset and Benchmark for Large Language Models for Protein Understanding (2024). URL https://arxiv.org/abs/2406.05540.

[32] Passaro, S. et al. Boltz-2: Towards Accurate and Efficient Binding Affinity Prediction (2025). URL 10.1101/2025.06.14.659707.

[33] Candido, S. et al. Language Modeling Materializes a World Model of Protein Biology (2026). URL 10.64898/2026.06.03.729735.

[34] Le Guilloux, V., Schmidtke, P. & Tuffery, P. Fpocket: An open source platform for ligand pocket detection. BMC Bioinformatics 10, 168 (2009). URL 10.1186/1471-2105-10-168.

[35] Trott, O. & Olson, A. J. AutoDock Vina: Improving the speed and accuracy of docking with a new scoring function, efficient optimization, and multithreading. Journal of Computational Chemistry 31, 455–461 (2010). URL 10.1002/jcc.21334.

[36] McNutt, A. T. et al. GNINA 1.0: molecular docking with deep learning. Journal of Cheminformatics 13, 43 (2021). URL 10.1186/s13321-021-00522-2.

[37] Eastman, P. et al. OpenMM 7: Rapid development of high performance algorithms for molecular dynamics. PLOS Computational Biology 13, e1005659 (2017). URL 10.1371/journal.pcbi.1005659.

[38] Baell, J. B. & Holloway, G. A. New Substructure Filters for Removal of Pan Assay Interference Compounds (PAINS) from Screening Libraries and for Their Exclusion in Bioassays. Journal of Medicinal Chemistry 53, 2719–2740 (2010). URL 10.1021/jm901137j.

[39] Daina, A., Michielin, O. & Zoete, V. SwissADME: a free web tool to evaluate pharmacokinetics, drug-likeness and medicinal chemistry friendliness of small molecules. Scientific Reports 7, 42717 (2017). URL 10.1038/srep42717.

