## Supplemental Materials for "A Persistent Fleet of AI Scientists Exhibits Cooperative and Autopoietic Behavior"

### Supplementary materials for

A Persistent Fleet of AI Scientists Exhibits Cooperative and Autopoietic Behavior

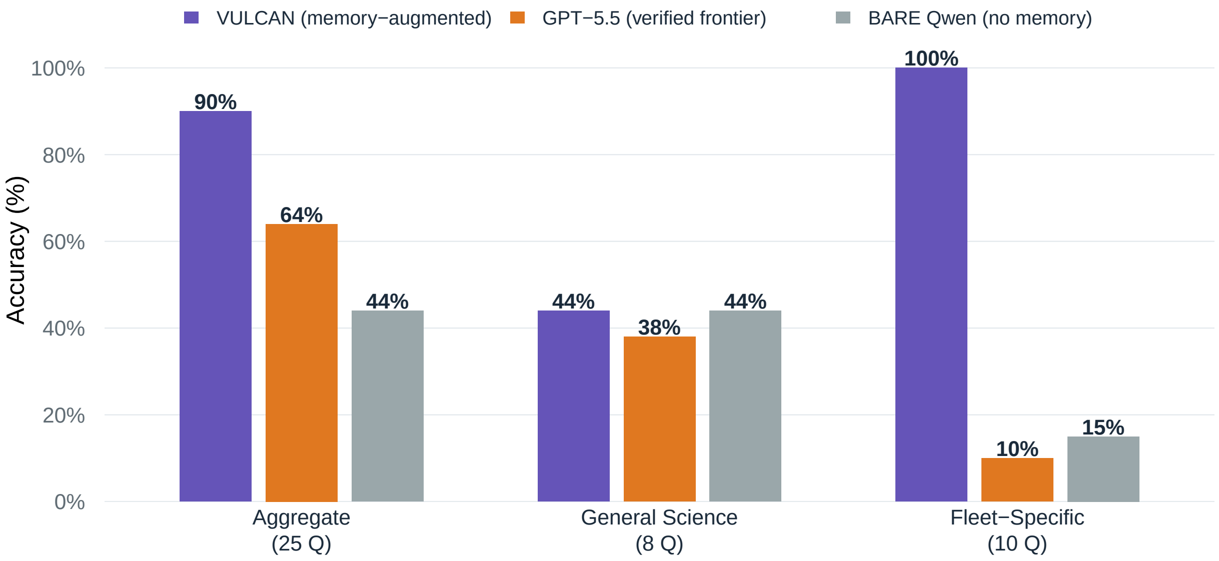


**Supplementary Figure S1. Memory functions as a lookup, not a reasoning upgrade** Triplicate, blind-scored 25-question benchmark comparing BARE Qwen, VULCAN memory-augmented Qwen, and GPT-5.5. VULCAN improves aggregate performance from 44% to about 90%, with the gain concentrated in fleet-specific questions that require access to local institutional knowledge. General-science performance is flat across conditions, supporting the interpretation that memory supplies specific information access rather than a generalized reasoning upgrade. The remaining 7/25 questions are included in the aggregate but are not disaggregated in the available category-level benchmark records, so the polished figure reports only confirmed category breakdowns.

**Supplementary Data S1 — VULCAN 25-question benchmark summary**

| Group | Denominator | VULCAN memory augmented (%) | GPT-5.5 frontier (%) | BARE Qwen no memory (%) | Note |
|---|---|---|---|---|---|
| Aggregate | 25 | 90 | 64 | 44 | Triplicate blind-scored aggregate; VULCAN 22.5/25, GPT-5.5 16/25, BARE 11/25 |
| General Science | 8 | 44 | 38 | 44 | No memory advantage on general science questions |
| Fleet-Specific | 10 | 100 | 10 | 15 | Memory provides access to fleet-specific facts |
| Other categories | 7 | not disaggregated | not disaggregated | not disaggregated | Aggregate scoring records confirm the total but not a per-category breakdown for the remaining 7 questions |

**Supplementary Table S2 — Released fleet skills**

| Skill | Category | Category count | Description | License/Release framing | Verification status | Evidence record |
|---|---|---|---|---|---|---|
| arxiv-fetch | Library Card Skills | 14 | Library card skill — programmatic access to arXiv preprints. Both metadata (Atom API) and PDFs are fully open: no Cloudf | MIT/public release subset per Vienna handoff; 20 UViiVe-built skills required MIT authorization at handoff time | Verified functional | UViiVE fleet-skills public release audit |
| biorxiv-fetch | Library Card Skills | 14 | Library card skill — programmatic access to bioRxiv/ medRxiv preprint METADATA (title, authors, abstract, DOI, date) via | MIT/public release subset per Vienna handoff; 20 UViiVe-built skills required MIT authorization at handoff time | Verified functional | UViiVE fleet-skills public release audit |
| semantic-scholar-api | Library Card Skills | 14 | Search academic papers, fetch citations/references, get author profiles, and retrieve paper details via the Semantic Sch | MIT/public release subset per Vienna handoff; 20 UViiVe-built skills required MIT authorization at handoff time | Verified functional | UViiVE fleet-skills public release audit |
| ncbi-api | Library Card Skills | 14 | Search PubMed, fetch abstracts, retrieve full-text links, and download open-access papers via NCBI E-utilities and PMC A | MIT/public release subset per Vienna handoff; 20 UViiVe-built skills required MIT authorization at handoff time | Verified functional | UViiVE fleet-skills public release audit |
| europepmc-search | Library Card Skills | 14 | Description not available in the public release audit. | MIT/public release subset per Vienna handoff; 20 UViiVe-built skills required MIT authorization at handoff time | Verified functional | UViiVE fleet-skills public release audit |
| abstract-lookup | Library Card Skills | 14 | Maximizes abstract coverage for research papers using a waterfall approach across multiple APIs (PubMed → Europe PMC → S | MIT/public release subset per Vienna handoff; 20 UViiVe-built skills required MIT authorization at handoff time | Verified functional | UViiVE fleet-skills public release audit |
| paperclip | Library Card Skills | 14 | Description not available in the public release audit. | MIT/public release subset per Vienna handoff; 20 UViiVe-built skills required MIT authorization at handoff time | Verified functional | UViiVE fleet-skills public release audit |
| openalex-search | Library Card Skills | 14 | Description not available in the public release audit. | MIT/public release subset per Vienna handoff; 20 UViiVe-built skills required MIT authorization at handoff time | Verified functional | UViiVE fleet-skills public release audit |
| chemrxiv-fetch | Library Card Skills | 14 | Library card skill — programmatic access to ChemRxiv chemistry preprints. Metadata via Crossref API (fully open). PDFs a | MIT/public release subset per Vienna handoff; 20 UViiVe-built skills required MIT authorization at handoff time | Verified functional | UViiVE fleet-skills public release audit |
| biorxiv-search | Library Card Skills | 14 | Description not available in the public release audit. | MIT/public release subset per Vienna handoff; 20 UViiVe-built skills required MIT authorization at handoff time | Verified functional | UViiVE fleet-skills public release audit |
| arxiv-search | Library Card Skills | 14 | Description not available in the public release audit. | MIT/public release subset per Vienna handoff; 20 UViiVe-built skills required MIT authorization at handoff time | Verified functional | UViiVE fleet-skills public release audit |

*Continued on next page*

**Supplementary Table S2 — Released fleet skills (continued)**

| Skill | Category | Category count | Description | License/Release framing | Verification status | Evidence record |
|---|---|---|---|---|---|---|
| pipette-skillgraph | Library Card Skills | 14 | Query the Pipette.bio Skill-Graph MCP server to discover bioinformatics workflows, pipeline transitions, and tool documen | MIT/public release subset per Vienna handoff; 20 UViiVe-built skills required MIT authorization at handoff time | Verified functional | UViiVE fleet-skills public release audit |
| summarize-pdf | Library Card Skills | 14 | Description not available in the public release audit. | MIT/public release subset per Vienna handoff; 20 UViiVe-built skills required MIT authorization at handoff time | Verified functional | UViiVE fleet-skills public release audit |
| docling-parse | Library Card Skills | 14 | Parse PDFs into structured markdown, JSON, or HTML using IBM Docling — handles tables, figures, multi-column layouts, an | MIT/public release subset per Vienna handoff; 20 UViiVe-built skills required MIT authorization at handoff time | Verified functional | UViiVE fleet-skills public release audit |
| pcr-primer-design | Design the Experiment | 5 | Design and validate primers for PCR, qPCR, TaqMan probes, multiplex, and sequencing assays with thermodynamic QC and MIQ | MIT/public release subset per Vienna handoff; 20 UViiVe-built skills required MIT authorization at handoff time | Verified functional | UViiVE fleet-skills public release audit |
| splicecraft | Design the Experiment | 5 | Agent-driven plasmid design, cloning simulation, and primer design via SpliceCraft local API. Supports Golden Braid/MoCl | MIT/public release subset per Vienna handoff; 20 UViiVe-built skills required MIT authorization at handoff time | Verified functional | UViiVE fleet-skills public release audit |
| experimental-design-statistics | Design the Experiment | 5 | Plan genomics experiments with power analysis, sample size estimation, batch-balanced layouts, and multiple-testing stra | MIT/public release subset per Vienna handoff; 20 UViiVe-built skills required MIT authorization at handoff time | Verified functional | UViiVE fleet-skills public release audit |
| genetic-variant-annotation | Design the Experiment | 5 | Annotate a VCF with functional consequences, clinical significance, population frequencies, and pathogenicity scores usi | MIT/public release subset per Vienna handoff; 20 UViiVe-built skills required MIT authorization at handoff time | Verified functional | UViiVE fleet-skills public release audit |
| gene-essentiality | Design the Experiment | 5 | Query, interpret, and integrate DepMap (Cancer Dependency Map) CRISPR knockout essentiality scores — including CERES, Ch | MIT/public release subset per Vienna handoff; 20 UViiVe-built skills required MIT authorization at handoff time | Verified functional | UViiVE fleet-skills public release audit |
| scrnaseq-scanpy-core-analysis | Analyze the Data | 8 | End-to-end scRNA-seq analysis in Python using Scanpy and the scverse stack — QC, normalization, integration, clustering, | MIT/public release subset per Vienna handoff; 20 UViiVe-built skills required MIT authorization at handoff time | Verified functional | UViiVE fleet-skills public release audit |
| scrnaseq-seurat-core-analysis | Analyze the Data | 8 | End-to-end scRNA-seq analysis in R using Seurat v5 — QC, SCTransform, Harmony / CCA / RPCA integration, clustering, anno | MIT/public release subset per Vienna handoff; 20 UViiVe-built skills required MIT authorization at handoff time | Verified functional | UViiVE fleet-skills public release audit |

*Continued on next page*

**Supplementary Table S2 — Released fleet skills (continued)**

| Skill | Category | Category count | Description | License/Release framing | Verification status | Evidence record |
|---|---|---|---|---|---|---|
| bulk-rnaseq-counts-to-de-deseq2 | Analyze the Data | 8 | End-to-end DESeq2 pipeline that turns a raw count matrix into a ranked, shrunken, QC-checked DE result table. Use when y | MIT/public release subset per Vienna handoff; 20 UViiVe-built skills required MIT authorization at handoff time | Verified functional | UViiVE fleet-skills public release audit |
| functional-enrichment-from-degs | Analyze the Data | 8 | Pathway / gene-set enrichment on differential-expression results using clusterProfiler. GSEA (rank-based, all genes) is | MIT/public release subset per Vienna handoff; 20 UViiVe-built skills required MIT authorization at handoff time | Verified functional | UViiVE fleet-skills public release audit |
| scrna-trajectory-inference | Analyze the Data | 8 | Reconstruct differentiation trajectories from scRNA-seq using PAGA, diffusion pseudotime, scVelo RNA velocity, and CellR | MIT/public release subset per Vienna handoff; 20 UViiVe-built skills required MIT authorization at handoff time | Verified functional | UViiVE fleet-skills public release audit |
| cell-cell-communication | Analyze the Data | 8 | Infer ligand-receptor signaling networks from annotated single-cell RNA-seq data and produce publication-ready communica | MIT/public release subset per Vienna handoff; 20 UViiVe-built skills required MIT authorization at handoff time | Verified functional | UViiVE fleet-skills public release audit |
| pooled-crispr-screens | Analyze the Data | 8 | Tiered analysis of pooled CRISPR screens with single-cell readout: fast t-test screening, on-target validation, and batc | MIT/public release subset per Vienna handoff; 20 UViiVe-built skills required MIT authorization at handoff time | Verified functional | UViiVE fleet-skills public release audit |
| spatial-transcriptomics | Analyze the Data | 8 | Run a complete 10x Visium pipeline — QC, clustering, spatially variable genes, neighborhood enrichment, co-occurrence — | MIT/public release subset per Vienna handoff; 20 UViiVe-built skills required MIT authorization at handoff time | Verified functional | UViiVE fleet-skills public release audit |
| data-analysis-best-practices | Report | 4 | Pre-analysis validation, missing-data handling, multiple-testing correction, batch-effect checks, reproducibility, and s | MIT/public release subset per Vienna handoff; 20 UViiVe-built skills required MIT authorization at handoff time | Verified functional | UViiVE fleet-skills public release audit |
| pdf-report-generation | Report | 4 | Generate professional, scientifically-styled PDF reports with ReportLab Platypus — title block, embedded figures, styled | MIT/public release subset per Vienna handoff; 20 UViiVe-built skills required MIT authorization at handoff time | Verified functional | UViiVE fleet-skills public release audit |
| docx-generation | Report | 4 | Generate professional, scientifically-styled Word (.docx) reports with python-docx — title block, tables, embedded figur | MIT/public release subset per Vienna handoff; 20 UViiVe-built skills required MIT authorization at handoff time | Verified functional | UViiVE fleet-skills public release audit |
| lasso-biomarker-panel | Report | 4 | Select a minimal (5–15 feature) biomarker panel from high-dimensional omics with nested-CV penalized logistic regression | MIT/public release subset per Vienna handoff; 20 UViiVe-built skills required MIT authorization at handoff time | Known disclosed Quick Start helper-script gap | UViiVE fleet-skills public release audit |

*Continued on next page*

**Supplementary Table S2 — Released fleet skills (continued)**

| Skill | Category | Category count | Description | License/Release framing | Verification status | Evidence record |
|---|---|---|---|---|---|---|
| skill-creator | Science That Thinks | 4 | Draft, validate, and package a new agentic-science skill — frontmatter, mandatory sections, reference docs, scripts, and | MIT/public release subset per Vienna handoff; 20 UViiVe-built skills required MIT authorization at handoff time | Verified functional | UViiVE fleet-skills public release audit |
| peer-review-gate | Science That Thinks | 4 | Description not available in the public release audit. | MIT/public release subset per Vienna handoff; 20 UViiVe-built skills required MIT authorization at handoff time | Verified functional | UViiVE fleet-skills public release audit |
| verified-science-review | Science That Thinks | 4 | High-confidence systematic literature review pipeline for biomedical research. Use when building comprehensive disease r | MIT/public release subset per Vienna handoff; 20 UViiVe-built skills required MIT authorization at handoff time | Verified functional | UViiVE fleet-skills public release audit |
| ensemble-analyze | Science That Thinks | 4 | Run a scientific question or dataset analysis through a configurable panel of frontier LLMs (Gemini, GPT-5.5, Claude Opu | MIT/public release subset per Vienna handoff; 20 UViiVe-built skills required MIT authorization at handoff time | Verified functional | UViiVE fleet-skills public release audit |

**Supplementary Data S2 — Benchmark summary and caveats**

| Claim area | Denominator/ design | Reported result | Evidence class | Required caveat | Evidence record |
|---|---|---|---|---|---|
| MIRROR First Law fabrication probe | 36-prompt main battery | Bare model passed 3/36 = 8.3%; First Law condition passed 36/36 = 100%; failure rate 91.7% to 0%; $P<0.001$ | Class I / published preprint | MIRROR preprint is canonical source; pre-publication May 1 data package is background only | MIRROR companion preprint and controlled benchmark record |
| VaaS-RIKER2 citation verification | 640 runs; C1 vs C3/C4 protocol comparison | Unguided Type II hallucination 95.9%; full protocol 6.5%; live-verification-only condition 0.0%; >14-fold reduction for the full protocol | Class I benchmark | C3 and C4 test distinct verification configurations; the manuscript reports C4 at 6.5% Type II and C3 at 0.0% | VaaS-RIKER2 640-run ablation benchmark record |
| Aletheia/Veraxis verification benchmark | 330 scored queries = 300-query seven-domain suite + earlier 30-query run | Sonar ~16% errors; calibrated Veraxis/Aletheia 0% emitted errors with HOLD/abstention | Pilot / AI-scored benchmark | Human validation pending; proxy/manual calibrated workflow, not final DrBlackwell clinical production gate | Aletheia/Veraxis 330-query audit record |
| VULCAN memory benchmark | 25 held-out questions, triplicate blind-scored | BARE 44%; VULCAN ~90%; GPT-5.5 64%; general-science category flat | Class I internal benchmark | Memory improves lookup and fleet-specific access, not general reasoning; 7/25 category details are not disaggregated in the available category-level benchmark records | Supplementary Figure S1 and Supplementary Data S1 |
| ProteinLMBench companion directionality | 944 independently authored literature-comprehension questions | Accuracy 54% without memory and 60% with memory | Class I external benchmark / internal memory toggle | The benchmark was independently authored; the memory-toggle evaluation was conducted by the fleet | ProteinLMBench publication and M4 memory-toggle result |
| Hypothesis Garden | 43 entries by cutoff; 14/23/6 distribution stated in manuscript | Class distribution supports Q3 patterning | Class II production records | One record with unresolved agent identities was excluded from decisive examples | Hypothesis Garden production corpus and classification record |
| Dream Cycle | 104 recurring cycles by reporting cutoff | Supports recursive, multi-checkpoint reasoning architecture | Class II production records | 104 is the manuscript reporting cutoff; subsequent cycles are outside the reported analysis | Nightly-cycle run history at the reporting cutoff |
| MEPAN/CFTR chaperone-design case | Two-stage computational personalized-medicine pipeline | G165Q established variant-specific stabilization logic; Y285C tested generalization; CFTR retrospective validated logic | Class III provenance case study | Computational/retrospective only; not wet-lab validation | MEPAN/CFTR computational case-study record |
| Dead-end registry scientific nulls | Production dead-end records | GP ensemble no improvement over baseline; pseudo-labeling does not fix OOD uncertainty | Class II production records | Production records are observational and were not independently blinded | Dead-end registry production audit record |
| 35 released skills / toolkit | 35 skills; 34/35 functional; 1 disclosed defect | Public subset organized into 14/5/8/4/4 categories | Toolkit / software supplement | lasso-biomarker-panel Quick Start helper-script gap disclosed | Supplementary Table S2 and fleet-skills public release audit |
